# Neonatal oxytocin prevents sex-specific spatial memory deficits induced by maternal separation through restoration of hippocampal synaptic plasticity in males

**DOI:** 10.64898/2026.07.04.736473

**Authors:** Hannah Illouz, Mila Jesic, Edith Tanche, Vincent Lelievre, Sylvain Hugel, Pierrick Poisbeau

## Abstract

**Background:** Stress during critical developmental periods causes lasting neurobiological alterations. Rodent models like neonatal maternal separation (NMS) induce cognitive alterations, particularly spatial memory deficits. Oxytocin (OT) system has been suggested to underlie these consequences, as it is critical for neurodevelopment. This neuropeptide also promotes maternal nurturing, prevents neuroinflammation and displays anxiolytic properties. This study hypothesized that early postnatal OT administration could prevent NMS-induced memory alterations in adult rats.

**Methods:** Sprague-Dawley rat pups (both sexes, n=8-12/group) underwent NMS with concomitant intraperitoneal OT injections. At adulthood, novel object recognition and object location tasks were performed. Further investigation was conducted through ex vivo electrophysiological recordings of functional plasticity at Schaffer collateral-CA1 synapses (male, n=7-12/group), alongside RT-qPCR of synaptic, GABAergic, neuro-inflammatory, and oxytocin receptor markers in dorsal CA1 (male, n=4-6/group).

**Results:** NMS induced male-specific spatial memory impairment without affecting recognition memory. Early OT completely prevented spatial memory deficits in NMS males. Electrophysiological recordings revealed that NMS suppressed CA1 long-term potentiation (LTP), and neonatal OT restored it. NMS induced transcript overexpression of neuro-inflammatory markers, GABAergic markers, and synaptic proteins in dorsal CA1. OT treatment normalized or reduced these mRNA expressions, consistent with restoration of CA1 synaptic function.

**Conclusion:** Early postnatal OT prevents NMS-induced spatial memory deficits and hippocampal LTP impairments in male rats, which is associated with normalized or reduced neuro-inflammatory and GABAergic transcript expressions. These findings establish exogenous oxytocin administration during a critical neonatal window as sufficient to prevent male-specific hippocampal dysfunction and cognitive deficits induced by early-life stress, identifying the oxytocinergic system as a promising target for early neuroprotective interventions.

## Introduction

The first thousand days of a child’s life represent a period of particularly high rates of synaptic regrowth and remodeling in the brain, during which environmental experiences are critical for brain development (Moore et al., 2017; Indrio et al., 2023). These experiences shape neural plasticity and influence children’s behavior and psychological processes throughout the lifespan, having long-lasting effects across a wide range of domains (Andersen, 2003; Pechtel and Pizzagalli, 2011; Melchior et al., 2022). Thus, chronic stress or trauma during early childhood, conceptualized as adverse childhood experiences (ACEs), have significant consequences for neurodevelopment, including multiple co-occurring mental and physical disorders (Felitti et al., 1998). More specifically, there is growing evidence that ACE-sensitive periods play a role in the development of neurobiological alterations, such as functional and volumetric changes in the amygdala and hippocampus, two key structures involved in emotion regulation and memory processing (Herzog and Schmahl, 2018). Rodent models of early life stress (ELS), such as neonatal maternal separation (NMS), represent useful paradigms to study the consequences of these ACEs. Behavioral and neuroendocrine signs of elevated stress reactivity have been observed in this model, as well as nociceptive and cognitive deficits, either immediately after stress exposure or as a long-term vulnerability in later life (Plotsky and Meaney, 1993; Ladd et al., 2000; Aisa et al., 2007; Melchior et al., 2022; Illouz et al., 2025).

The hippocampus is one of the primary targets of ELS as this brain area is undergoing major plastic changes during the first postnatal weeks (Rice and Barone, 2000). This structure presents a high expression of adrenal steroid receptors, which makes it particularly vulnerable to early environmental factors like stress or immune activation, alongside its high plasticity (McEwen, 1999, 2001; Hoeijmakers et al., 2015). After maturation, this key region enables stress regulation and emotional processing as part of the broader limbic system (McEwen, 1999, 2001). The hippocampus is also known to regulate episodic, declarative and spatial memory via synaptic plasticity processes between hippocampal subregions (Martin et al., 2000; Broadbent et al., 2004; Insausti and Amaral, 2004). The most intensively studied hippocampal pathway is the Schaeffer collateral projections from *cornu ammonis* 3 (CA3) pyramidal cells to CA1, a subregion enabling integration of novelty thanks to its large network of place cells (Sosa et al., 2016). Changes in either basal transmission or synaptic plasticity (long-term potentiation (LTP) and long-term depression (LTD) mechanisms) are believed to drive the behavioral consequences of ELS and are therefore important to investigate. Indeed, key structural synaptic proteins such as the presynaptic vesicular protein synaptophysin and the postsynaptic scaffolding protein PSD95, show reduced hippocampal expression following NMS (Wang et al., 2020). ELS has also been shown to alter functional plasticity mechanisms in the CA1 area of the hippocampus and in the dentate gyrus (DG) in a sex specific manner: male rodents present LTP impairment after chronic stress paradigms like NMS, while female remain unaffected (Derks et al., 2016, 2017). These molecular and cellular consequences are observed at the behavioral level, as memory tasks are poorly performed by rodents having experienced ELS (Alves et al., 2022). Recently, NMS male rats (but not female) were proved to display spatial memory deficits associated with an intense neuroinflammatory signature specifically in the dorsal CA1 (Illouz et al., 2025). Early and long-lasting neuroinflammation has also been previously proposed to sensitize the developing hippocampus, creating vulnerability for cognitive deficits following ELS (Hoeijmakers et al., 2015; Catale et al., 2022; Lumertz et al., 2022). In fact, most studies on males exposed to ELS report short- and long-term memory deficits, while most studies on females show no significant effects (Alves et al., 2022). The combination of these results reveals a marked sexual dimorphism, with males being particularly vulnerable to hippocampal and neuroinflammatory alterations. The mechanisms underlying this sex-specific vulnerability, however, remain poorly understood.

Oxytocin (OT) is a nonapeptide synthesized by hypothalamic neurons and known for its key regulatory role and neurohormonal effects in development. By activating G protein-coupled OT receptors (OTR) in peripheral tissues, OT plays multiple roles during parturition (uterine contraction) and after delivery (milk ejection). OT is also synaptically released in the brain from parvocellular projections and by magnocellular hormone-like release (Ludwig and Leng, 2006). Oxytocin receptors are expressed in both excitatory pyramidal neurons and inhibitory interneurons throughout all subregions of the hippocampus, particularly CA3 and CA2, though a weaker expression is also observed in DG and CA1 (Cilz et al., 2019; Young and Song, 2020). While systemic OT has been reported to promote facial recognition and maternal nurturing, to prevent neuroinflammation and to display anxiolytic as well as analgesic properties (Melchior et al., 2018; Mairesse et al., 2019), hippocampal OT is specifically considered to modulate several aspects of learning, memory and sociability (Tomizawa et al., 2003; Raam et al., 2017), as well as neuronal activity (Cilz et al., 2019; Zagrean et al., 2022). Perinatal oxytocin receptor (OTR) signaling in hippocampal neurons is neuroprotective, and shapes circuit development in young offspring (Cilz et al., 2019; Wu et al., 2021). It notably reduces deleterious stress effects on hippocampal synaptic plasticity and memory in rats thanks to its regulation of ERK activity (Lee et al., 2015). Transient OT may also control hippocampal glutamatergic and GABAergic circuit development, and contribute to the maintenance of an optimal physiological excitation/inhibition (E/I) ratio (Ripamonti et al., 2017). In fact, early postnatal OT signaling enables maturation of inhibitory GABAergic signaling through upregulation of KCC2 expression, which lowers intracellular chloride concentration and initiates GABAA receptor-mediated hyperpolarizing and inhibitory effects (Leonzino et al., 2016).

NMS has been repeatedly associated with oxytocinergic signaling dysregulation (Melchior et al., 2018; Baracz et al., 2020), as well as with an impaired GABA maturation leading to an imbalance between excitation and inhibition, influencing long-term neurodevelopment (Gazzo et al., 2021; Illouz et al., 2025). Given the critical roles of OT in neurodevelopment, the hippocampal vulnerability to early-life stress, and the sex-specific nature of NMS-induced alterations, we hypothesized that early postnatal OT administration could prevent the development of memory deficits and associated neurophysiological changes in NMS male rats.

In this study, we administered exogenous OT by means of intraperitoneal injection to NMS offspring to assess its possible preventive effects. We characterized long-term behavioral consequences in both male and female rats using short-term novel object recognition (recognition memory) and object location tasks (spatial memory). Given the male-specific deficit observed in spatial memory, we further investigated the underlying mechanisms exclusively in males through *ex vivo* electrophysiological recordings of LTP and LTD at Schaffer’s collateral synapses in dorsal hippocampus. This was accompanied by a molecular analysis of synaptic proteins (synaptophysin, PSD95), GABAergic (GAD65, NKCC1, KCC2), and neuroinflammatory (CD11b, GFAP, TNFα) markers, as well as oxytocin receptor expression in the dorsal CA1 region.

## Materials and Methods

### Animals

Pregnant female Sprague-Dawley rats (Charles River, Saint-Germain Nuelles, France) were used in this study and carefully monitored for delivery. The day of birth was referred to as postnatal day 0 (P0). Mother and pups were housed in a temperature (22±2 ◦C) and humidity (45 ± 10%) controlled room, under a 12-hour light-dark cycle (lights on at 7:00 am), with ad libitum access to food and tap water. Pups were weaned at P21 and housed in collective cages according to sex. Males and females were initially used in this study, then only males for RT-qPCR and electrophysiology experiments, in accordance with male-specific differences observed during behavioral tests. Different litters were used to perform behavioral tests (3 litters/group), as well as molecular (2 litters/group) and electrophysiological analyses (3 litters/group). All procedures were conducted in accordance with EU regulations and approved by the regional ethical committee (CREMEAS authorization numbers APAFIS#41332-2023021317122829 v6).

### Neonatal Maternal Separation

Litters were randomized at birth into two groups: nonseparated control litters and neonatal maternal separated ones (NMS). From postnatal days 2 (P2) to 12 (P12), the litters assigned to the NMS group were taken out of the nest cages three hours a day and put in a different cage under a heating lamp (28±2°C, wavelengths between 1400nm and 3000nm). During separation time, litters assigned to the control group stayed in their home cages with their mothers and did not get any additional treatment aside from changing their cage bedding, and vehicle injections. Before beginning behavioral tests, the pups were weaned at P21 and kept in cages with four rats each.

### Drug Treatments

All molecules used in this study were injected intraperitoneally (i.p. 10 µl, 30-gauge needle) daily during the NMS period, i.e., from P2 to P12 (11 injections in total). The injections were performed at the beginning of the separation process.

NaCl (0.9%) was used as a vehicle to control the sole effect of the neonatal injections. Oxytocin (OT, Bachem, Weil am Rhein, Germany) was diluted 5min before the experiment to inject a daily final dose of 1mg/kg to the pups in the NMS groups, as described previously (Melchior et al., 2018).

### Behavioral Testing

Animals were tested between P45 and P55 and chosen randomly in at least three different litters. Experiments were performed during the light phase, between 9:00a.m. and 6:00p.m. Prior to behavioral testing, animals were habituated to the experimenter with one week of handling.

### NORT and OLT

The object recognition test was adapted from Ennaceur and Delacour (Ennaceur and Delacour, 1988). The open field was composed of black wood and was 65 cm by 65 cm by 45 cm. The animals were acquainted with the field for fifteen minutes on the three days before trial. In the first attempt of the experiment, two similar objects were put inside the chamber at equal distances (10 cm) from the sides (2 rectangular plastic boxes, h 40cm x L 15cm x l 5cm). Prior to testing, all objects were assessed for baseline preferences to ensure no inherent biases existed between objects. Rats were placed with their nose facing the wall in a corner of the open field and allowed to freely explore for 15 minutes. One hour later, a second trial took place, in which one object was replaced by a different one (1 round iron box, d 5cm x h 35cm), and exploration was scored for 4 minutes. To avoid preference for one of the objects, the order of the objects was balanced between the testing animals.

One week later, the same scheme was used for the object location test (three days habituation sessions, one training phase and a 1-h testing trial), now using two identical objects but changing their location (2 glass bottles, d 10cm x h 30cm). Rats were always placed with their nose facing the wall, like during training phase. One hour after training, one of the objects was relocated in the diagonal corner relative to the other object. In the same way as for NORT, which of the two objects (the left or right object) was relocated, as well as the wall upon which it was organized, was counterbalanced.

Each test phase (habituation, training phase, and testing trial) was filmed to be analyzed using a camera, and exploration of the two objects was scored for 4 minutes. During analysis, it was considered that the animal was exploring the object when the head of the rat was oriented toward the object with its nose within 2 cm of the object. Results were expressed as the difference in the exploration time of the two objects divided by the total time spent exploring the objects (discrimination ratio) (Ennaceur and Delacour, 1988; Mendez et al., 2015). In order to eliminate olfactory stimuli, chambers and objects were always cleaned after testing each animal.

### Molecular Analyses

Hippocampal tissues were collected from P60 rats in order to analyze the dorsal CA1 of the hippocampus individually (Figure 1A). The brains were harvested, stored at -80°C, and then sliced into 250µm slices with a cryotome (Leica CM3050 S Cryostat). Small needles (Fine Science Tools, item 18035-80; length 8.5 cm, diameter 0.8 mm) were used to harvest tissue disks (by puncture) from the dorsal CA1, CA3, and dentate gyrus regions of each animal’s hippocampus.

**Figure 1:**
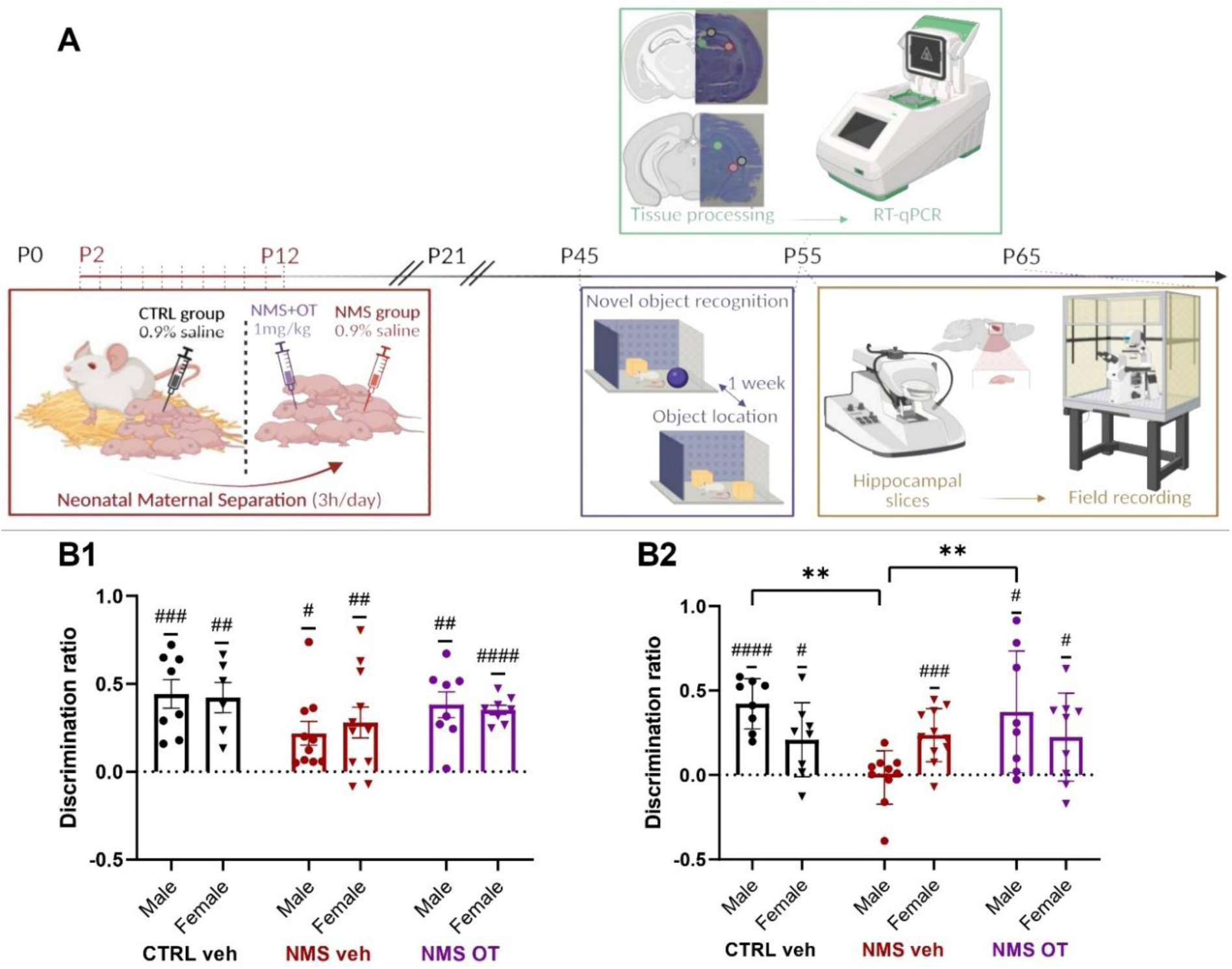
NMS induces sex-specific spatial memory alterations prevented by neonatal oxytocin. **A:** After birth (P0), randomly selected litters underwent daily injection of vehicle or OT before undergoing NMS or not. All pups remained with their mothers until weaning at P21. Mnesic testing was conducted between P45 and P55, followed by RT-qPCR or electrophysiological analyses in the dorsal CA1 hippocampal regions. **B1:** No alterations were observed between groups concerning recognition memory. **B2:** NMS+veh rats showed a male-specific spatial memory deficit. This alteration was prevented by early OT injections. Statistical significance was assessed with one sample t test against 0, and illustrated as follows: p<0.05 (#), p < 0.01 (##), p < 0.001 (###) or p < 0.0001 (####). Sex and group effects were assessed using 2-way ANOVA followed by Tukey’s multiple comparison post-hoc test and illustrated as follows: p < 0.01 (**). Male CTRL+veh: n=8, Female CTRL+veh: n=8, Male NMS+veh: n=10, Female NMS+veh: n=11, Male NMS+OT: n=8, Female NMS+OT: n=8.

### Quantitative RT-qPCR

Total RNA was extracted using a protocol adapted from the original procedure of Chomczynski and Sacchi (Chomczynski and Sacchi, 1987) consisting in 2 independent total RNA extractions separated by a DNAse I treatment (TURBOTM DNase; Ambion, Life technologies, Saint Aubin, France), as previously described in detail (Lelievre et al., 2002). 600 ng RNA were reverse transcripted with the RT iScript kit (Bio-Rad, Marnes-la-Coquette, France). Quantitative PCR was performed using SYBR Green Supermix (Bio-Rad), on the iQ5 Real Time PCR System (Bio-Rad). Amplifications were carried out in 45 cycles (20 s at 95 ◦C, 20 s at 60 ◦C, and 20 s at 72 ◦C). Primer sets for all genes of interest were designed using Oligo6.0 and M fold softwares (primer sequences in Supplementary Table 1, also used in Illouz *et al*., 2025. A melt curve was run for each primer to verify the formation of only one product. Data were normalized to the housekeeping GAPDH (Glyceraldehyde 3-phosphate dehydrogenase) using the ΔΔct method (Livak and Schmittgen, 2001). Statistical analyses validated GAPDH stability between experimental groups.

### Hippocampal Electrophysiology

#### Acute Slice Preparation

P55 to P65 rats were deeply anesthetized with an intraperitoneal injection of ketamine/xylazine (180/20 mg/kg). Under deep anesthesia, intracardiac perfusion was performed using oxygenated, ice-cold (0–4 °C) sucrose-based artificial cerebrospinal fluid (sACSF), continuously bubbled with carbogen (95% O₂, 5% CO₂). The sACSF solution contained (in mM): 248 sucrose, 11 glucose, 26 NaHCO₃, 2 KCl, 1.25 KH₂PO₄, 2 CaCl₂, 1.3 MgSO₄, and 2.5 kynurenic acid.

After approximately 15 minutes of perfusion, the animals were decapitated, and their brains were rapidly removed. Using a brain matrix and razor blade, the anterior region (rostral to the corpus callosum) and the posterior region (at the level of the cerebellum) were excised. The remaining tissue was mounted onto the stage of a vibratome (VT1200S; Leica, Nussloch, Germany) by gluing the posterior end.

Transverse dorsal hippocampal slices (450 µm thick) were prepared and immediately transferred to a chamber containing standard artificial cerebrospinal fluid (aCSF), maintained at room temperature. The aCSF composition was (in mM): 126 NaCl, 26 NaHCO₃, 2.5 KCl, 1.25 NaH₂PO₄, 2 CaCl₂, 2 MgCl₂, and 10 glucose, bubbled continuously with 95% O₂ and 5% CO₂ (pH 7.3; ∼310 mOsm, measured).

#### Field Potential Recordings

After at least 1 hour of recovery, individual hippocampal slices were transferred to a submerged recording chamber and continuously perfused with oxygenated aCSF at a rate of 2.5–3 ml/min. Extracellular field excitatory postsynaptic potentials (fEPSPs) were recorded in the *stratum radiatum* of the CA1 region following stimulation of the Schaffer collateral pathway.

Stimulation and recording electrodes were fabricated from borosilicate glass capillaries (1.2 mm inner diameter, 1.69 mm outer diameter; Warner Instruments, Harvard Apparatus) using a P-1000 horizontal puller (Sutter Instruments). Both electrodes had a resistance of approximately 3 MΩ when filled with aCSF. Recordings were performed using a MultiClamp 700A amplifier (Molecular Devices). Signals were sampled at 20 kHz, digitized with a BNC-2110 data acquisition interface (National Instruments), and acquired using WinWCP software (Strathclyde Electrophysiology Software, John Dempster, University of Strathclyde, UK).

To determine stimulus intensity, we gradually increased stimulation amplitude until the fEPSP response reached a maximum, then reduced it to evoke responses at approximately two-thirds of the maximal amplitude. Test stimuli were delivered at a frequency of 0.033 Hz (one pulse every 30 seconds).

#### Plasticity Protocols

Baseline synaptic transmission was monitored for at least 15 minutes prior to induction protocols. Long-term synaptic plasticity was induced using either high frequency stimulation (HFS) or low-frequency stimulation (LFS). HFS consisted of trains of five pulses at 100 Hz, repeated six times at 10-second intervals. LFS consisted of 900 pulses delivered at 1 Hz for 15 minutes. Synaptic responses were monitored for 60 minutes following induction.

#### Statistical Analyses

For each experiment, normality of residuals and homoscedasticity was verified with Shapiro-Wilk and Spearman’s tests prior to performing parametric analysis. Differences were considered statistically significant at p < 0.05. Data are expressed as mean ± standard error of the mean (SEM).

Concerning memory tests, the novel object recognition and object location tasks were analyzed with one sample t test with 0 as theoretical mean, 0 being the value for which the animals have no preference for any of the objects. Inter- and intra-group effects were assessed using 2-way ANOVA (group x sex) followed by Tukey’s multiple comparison post-hoc test. Statistical analysis was performed using GraphPad Prism 9 software (La Jolla, USA).

fEPSP slopes were measured using Clampfit 11.4.38. For each recording, slopes were normalized to the mean baseline slope recorded during the 10 min pre-induction period. Analyses following induction were limited to the 5–60 min period, as responses during the first 0–5 min were affected by post-tetanic potentiation. fEPSPs were binned into 5-min non-overlapping intervals, and log-transformed normalized fEPSP slopes were used for statistical analyses. Block averages were calculated for 5–20 min, 20–40 min, and 40–60 min intervals. The baseline period was included to allow within-treatment comparisons (post-induction vs baseline). Statistical comparisons were performed with a global window (5–60 min) and block-wise analyses (5–20, 20–40, 40–60 min). Paired Wilcoxon signed-rank tests were used to compare each recording’s to its baseline. Intergroup comparisons were performed for each block with Kruskal–Wallis one-way ANOVA (1wANOVA) to detect overall differences. Mann–Whitney U tests were used for pairwise post-hoc comparisons. All electrophysiological analyses were performed using Python (pandas, numpy, scipy).

All PCR results were processed using the ΔΔct method (Livak and Schmittgen, 2001) conducted in R studio (version 4.5.1; readxl, dplyr, writexl). Intergroup comparisons of transcript expressions were done using 1wANOVA followed by Tukey’s multiple comparison post-hoc test (GraphPad Prism 9 software, La Jolla, USA).

## Results

Cognitive long-term effects of NMS and dOVT injections were characterized at young adulthood, between P45 and P55 (fig.1A).

The novel object recognition test showed that CTRL+veh, NMS+veh and NMS+OT rats were able to discriminate between a previously encountered object and a novel object, as the mean discrimination ratios were significantly higher than identical exploration of the two objects (Fig. 2B1; unpaired sample t tests ; CTRL+veh M: P=0.0009; df=7 – CTRL+veh F: P=0.0044; df=5 – NMS+veh M: P=0.0109; df=9 – NMS+veh F: P=0.0099; df=10 – NMS+OT M: P=0.0013; df=7 – NMS+OT F: P<0.0001; df=7). No intergroup differences were observed (2wANOVA, group: F (2,45) = 3.086, p=0.0555).

**Figure 2:**
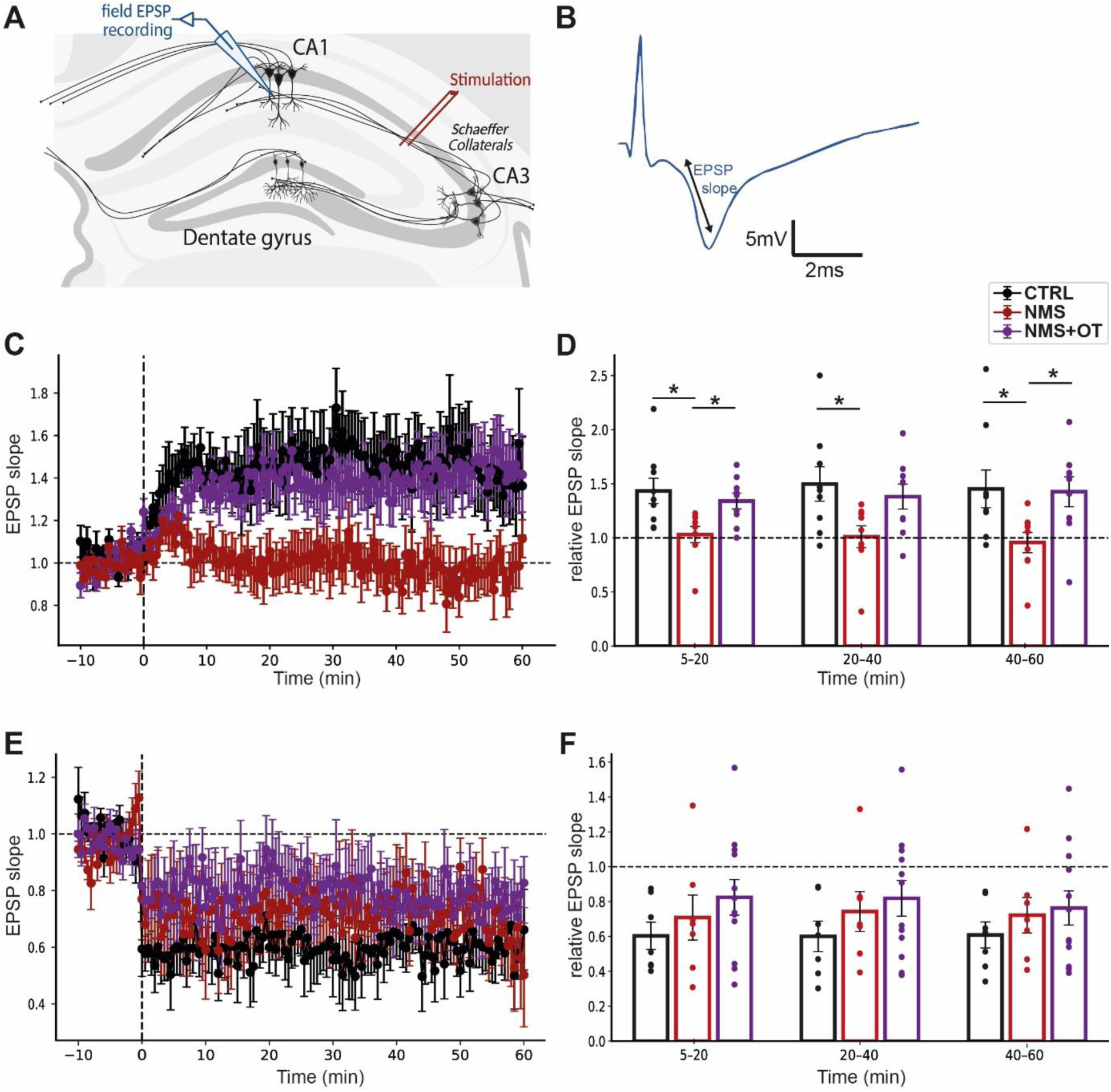
NMS induces long-term plasticity alterations in adult male rats prevented by neonatal oxytocin. **A:** Schematic representation of the placement of stimulating and recording electrode in the Schaffer collateral-CA1 pathway in the hippocampus. **B:** Representative signal trace recorded from stratum radiatum of CA1. **C:** Time course plots of normalized postsynaptic potentials (EPSP) recorded before and after HFS of the Schaffer collateral-CA1 pathway. CTRL+veh and NMS+OT groups, but not NMS+veh, were associated with LTP increases, as also represented on the corresponding histogram representing average potentiation after HFS (**D**). **E:** Time course plots of normalized EPSP recorded before and after LFS of the Schaffer collateral-CA1 pathway. All groups presented an intact LTD, as indicated on the bar histogram depicting average depression after HFS (**F**). LTP and LTD induction protocols were applied at t=0. Statistical significance was assessed with paired Wilcoxon signed-rank tests to compare each recording’s mean log-transformed normalized fEPSP slope to baseline. Group effects were assessed using Kruskal-Wallis 1-way ANOVA followed by Mann-Whitney U’s multiple comparison post-hoc test and illustrated as follows: p < 0.05 (*). LTP: CTRL+veh: n=9, NMS+veh: n=9, NMS+OT: n=9, LTD: CTRL+veh: n=7, NMS+veh: n=7, NMS+OT: n=12.

In the object location test, male and female CTRL+veh rats recognized the moved object (mean discrimination ratios higher than zero) (Fig. 2B2; unpaired sample t tests ; CTRL+veh M: P<0.0001; df=7 – CTRL+veh F: P=0.0313; df=7). The same applies for female from the NMS+veh group (unpaired sample t test ; NMS+veh F: P=0.0005; df=10). However, NMS male with early vehicle injections have difficulties to discriminate between a previously encountered object location and a novel location (Fig. 2B2; unpaired sample t test ; NMS+veh M: P=0.7752; df=9). Male and female NMS+OT recognized the moved object, suggesting that early postnatal OT injections restored spatial memory (Fig. 2B2; unpaired sample t test ; NMS+OT M: P=0.0220; df=7 – NMS+OT F: P=0.0323; df=8). A group and interaction effect was highlighted, with a specific difference between male CTRL+veh and male NMS+veh, and between male NMS+veh and male NMS+OT rats (2wANOVA, interaction effect: F(2,48) = 6.002, P=0.0047 – Tukey’s multiple comparison test, CTRL+veh M vs NMS+veh M, P=0.0021, NMS+veh M vs NMS+OT M, P=0.0081).

The consequences of NMS on plasticity mechanisms in the dorsal hippocampal CA1 in adult males were then assessed, as well as whether daily neonatal OT injections would prevent potential LTP or LTD alterations.

High-frequency stimulation induced lasting LTP in CTRL+veh male rats, with normalized fEPSP slopes significantly higher than baseline. By contrast, high-frequency stimulation failed to induce fEPSP potentiation in NMS+veh animals, reflecting impaired hippocampal plasticity. Interestingly, rats treated with OT between P2 and P12 showed significant LTP in adulthood, as their responses exceeded baseline levels (Fig. 2C; log-transformed normalized means; paired Wilcoxon signed-rank test: CTRL+veh: P=0.0078; df=8 – NMS+veh : P=0.6523; df=8 – NMS+OT : P=0.0273; df=8). Initiation of LTP (first 20 minutes post-HFS) was significantly stronger in CTRL+veh and NMS+OT groups compared with NMS+veh animals (Fig. 2D; Kruskal-Wallis 1w ANOVA, χ²(2), P=0.0154 – Mann-Whitney multiple comparison test, CTRL+veh vs NMS+veh, P=0.0142, NMS+veh vs NMS+OT, P=0.0106). This pattern persisted for late-LTP (40–60 minutes post-HFS), with both CTRL+veh and NMS+OT maintaining potentiation well above that of NMS+veh (Kruskal-Wallis 1w ANOVA, χ²(2), P=0.0123 – Mann-Whitney multiple comparison test, CTRL+veh vs NMS+veh, P=0.0142, NMS+veh vs NMS+OT, P=0.0078). Between 20 and 40 minutes, the same trend is visually observed, i.e., high potentiation only for the CTRL+veh and NMS+OT groups (Kruskal-Wallis 1w ANOVA, χ²(2), P=0.0651 – Mann-Whitney multiple comparison test, CTRL+veh vs NMS+veh, adjusted P=0.0244). These results demonstrate that neonatal OT prevented impairments in adult LTP, both in terms of amplitude and maintenance.

In contrast, low-frequency stimulation at the Schaeffer collateral induced LTD in each of the three groups studied, with no detectable impact of NMS or neonatal OT injections on this specific type of plasticity. Thus, fEPSP slopes were significantly lower than baseline in each group (Fig. 2E; paired Wilcoxon signed-rank test; CTRL+veh: P=0.0156; df=6 – NMS+veh : P=0.0469; df=6 – NMS+OT : P=0.0425; df=11). No intergroup differences were observed in any of the periods analyzed (Fig. 2F; Kruskal-Wallis 1w ANOVA, χ²(2), 5-20min: P=0.3892, 20-40min: P=0.3844, 40-60min: P=0.7684).

NMS has been shown to induce a neuroinflammatory response in the dorsal CA1 hippocampus of male rats (Illouz et al., 2025), as well as male- and dorsal CA1-specific changes in oxytocinergic signaling and chloride cotransporters, suggesting plastic mechanisms affecting local neuronal inhibition. We therefore measured the expression of these markers, as well as synaptic markers, in order to determine whether the restoration of a behavioral and functional phenotype similar to CTRL+veh observed in NMS+OT animals is also detectable at the molecular level. Figure 3A recapitulates mRNA overexpression in NMS+veh groups compared to CTRL+veh animals, regarding the neuroinflammatory, neuronal inhibition, synaptic and oxytocinergic markers. NMS+OT male rats appear to have similar (CD11b, TNFα, OTR) or reduced (GFAP, GAD65, NKCC1, KCC2, SYN, PSD95) transcript expression in comparison to CTRL+veh rats, suggesting a powerful restorative effect of neonatal OT at the molecular level, which could even reverse the expression of certain transcript at adulthood.

**Figure 3:**
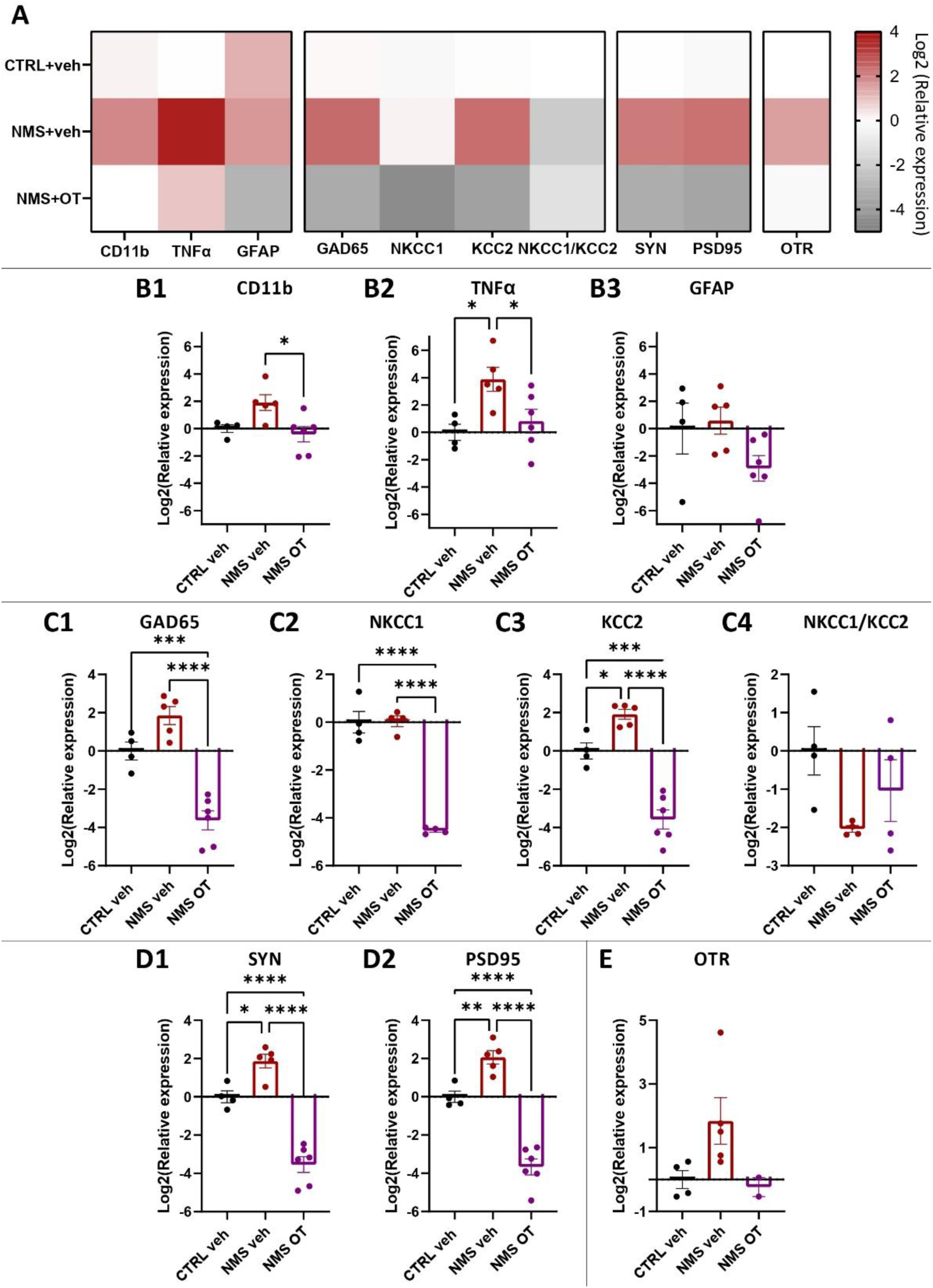
Changes in relative expression of dorsal CA1 hippocampal transcripts in CTRL+veh, NMS+veh and NMS+OT male rats (P60) as measured with RT-qPCR. **A:** NMS+veh group was associated with transcript increases concerning neuroinflammatory (**B**), inhibition (**C**), synaptic (**D**) and oxytocinergic markers (**E**), while early OT injections seem to inverse these consequences (no difference with CTRL group or inverse effect of NMS+veh). Statistical significance between groups was assessed with a 1wANOVA followed by Tukey’s multiple comparison post-hoc test, illustrated as follows: p < 0.05 (*), p < 0.01 (**), p<0.0001 (****). CTRL+veh: n=4, NMS+veh: n=5, NMS+OT: n=6.

In accordance with these visual results, intergroup comparison points out a 3.7 mRNA overexpression of the microglial marker CD11b (fig. 3B1) and a 14.7 times higher transcript expression of the proinflammatory cytokine TNFα (fig. 3B2) in the dCA1 of NMS+veh animals compared to CTRL+veh rats. Interestingly, NMS+OT male rats show a return to basal expression of these neuroinflammatory transcripts (1wANOVA test followed by Tukey’s multiple comparison tests ; <u>CD11b:</u> P=0.019, F (2, 12)=5.611, CTRL+veh vs. NMS+veh P=0.0819, NMS+veh vs. NMS+OT P=0.0186 – <u>TNFα:</u> P=0.0177, F (2, 12)=5.75, CTRL+veh vs. NMS+veh P=0.0229, NMS+veh vs. NMS+OT P=0.0462). However, GFAP (fig. 3B3) mRNA expression did not differ between CTRL+veh, NMS+veh and NMS+OT groups, even though this group shows a 7.5 decrease visually (1wANOVA test ; P=0.1126, F (2, 12)=2.634).

The same pattern as CD11b and TNFα applies for markers of neuronal inhibition (fig. 3C), with overexpression of GAD65 and KCC2 transcripts in NMS+veh animals alongside underexpression of GAD65 (fig. 3C1), NKCC1 (fig. 3C2), and KCC2 (fig. 3C3) in the NMS+OT group (1wANOVA test followed by Tukey’s multiple comparison tests ; <u>GAD65:</u> P<0.0001, F (2, 12)=35.28, CTRL+veh vs NMS+veh P=0.0665, CTRL+veh vs. NMS+OT P=0.0007, NMS+veh vs. NMS+OT P<0.0001 – <u>NKCC1:</u> P<0.0001, F (2, 9)=80.5, CTRL+veh vs. NMS+OT P<0.0001, NMS+veh vs. NMS+OT P<0.0001 – <u>KCC2:</u> P<0.0001, F (2, 12)=47.33, CTRL+veh vs NMS+veh P=0.0281, CTRL+veh vs. NMS+OT P=0.0002, NMS+veh vs. NMS+OT P<0.0001). This seem to result in disrupted NKCC1/KCC2 ratios (fig. 3C4), representing a deregulation of the excitation-inhibition balance in favor of inhibition. Even if this result isn’t statistically significant, chloride extrusion could be 4 times increased in the NMS+veh group, and half as much in the NMS+OT group, highlighting a tendency to return to a basal balance in the NMS+OT group (Brown-Forsythe ANOVA test ; P=0.1311, F* (2, 5.756) = 2.953).

With regard to the presynaptic marker SYN, the same pattern is present (3.6 times transcript overexpression in the NMS+veh group, 11.6 times under-expression in the NMS+OT group) (fig. 3D1; 1wANOVA test followed by Tukey’s multiple comparison tests ; P<0.0001, F (2, 12)=56.91 ; CTRL+veh vs NMS+veh P=0.0176, CTRL+veh vs. NMS+OT P<0.0001, NMS+veh vs. NMS+OT P<0.0001). The same applies to the postsynaptic marker PSD95 (fig. 3D2; 1wANOVA test followed by Tukey’s multiple comparison tests ; P<0.0001, F (2, 12)=63.56 ; CTRL+veh vs NMS+veh P=0.0094, CTRL+veh vs. NMS+OT P<0.0001, NMS+veh vs. NMS+OT P<0.0001).

Fig. 3E shows a visual 3.5 overexpression of NMS+veh OTR transcripts, alongside return to basal expression in NMS+OT rats (1wANOVA test: P=0.0834, F (2, 8) = 3.444).

## Discussion

The present study demonstrates that neonatal oxytocin administration during NMS prevents the emergence of male-specific spatial memory deficits and associated hippocampal dysfunction in adulthood. While NMS selectively impaired spatial memory in males, daily neonatal OT injections fully prevented this deficit. Electrophysiological recordings revealed that NMS abolished LTP at Schaffer collateral-CA1 synapses in males without impacting LTD. Early OT restored normal potentiation across both early induction and late maintenance phases. At the molecular level, NMS was associated with marked overexpression of neuroinflammatory transcripts (CD11b, TNFα), GABAergic developmental markers (GAD65, KCC2), and synaptic protein mRNAs (SYN, PSD95) in the dorsal CA1 region of males, all of which were normalized or reversed by early OT treatment. Together, these findings suggest that OT is a critical neuroprotective factor during early postnatal development that supports hippocampal maturation and prevents stress-induced cognitive dysfunction.

The sex-specific NMS alterations observed here converge with evidence for male-specific cognitive impairments to early-life adversity. NMS impaired hippocampus-dependent memory in males, but not females, and spared recognition memory (a perirhinal cortex-dependent process) (Winters et al., 2004). This replicates and extends recent reports demonstrating male-specific impairments in hippocampus-dependent memory tasks after early stress (Loi et al., 2017; Alves et al., 2022; Illouz et al., 2025). Consistent with this behavioral deficit, NMS selectively impaired LTP in the CA1 region of male rodents, but not females, particularly from 8 weeks onwards (Herpfer et al., 2012; Sousa et al., 2014; Derks et al., 2016, 2017). Previous results also reported that NMS male rats display spatial memory deficits associated with an intense neuroinflammatory signature specifically in dorsal CA1, while females remain unaffected (Illouz et al., 2025). This hippocampal vulnerability likely reflects the high density of glucocorticoid receptors in this region (Jacobson and Sapolsky, 1991; Lupien et al., 2009), combined with sustained hypothalamo-pituitary-adrenal (HPA) axis hyperreactivity and elevated corticosterone levels in adulthood following NMS (Plotsky and Meaney, 1993; Alves et al., 2020; Nishi, 2020). Whether OT exerts its neuroprotective effects partly through normalization of HPA reactivity stays an open question that future studies should address. The mechanisms underlying female resilience to NMS remain to be fully elucidated, and could involve sexually dimorphic developmental processes including estrogen-mediated OTR upregulation and differential microglial reactivity (Acevedo-Rodriguez et al., 2015; Breach and Lenz, 2023; Illouz et al., 2025).

The neuroinflammatory signature observed in NMS males (upregulation of CD11b and TNFα, marker of microglial activation and proinflammatory cytokine, respectively) is consistent with the emerging concept that ELS primes or sensitizes the developing hippocampus through sustained neuroimmune activation (Hoeijmakers et al., 2015; Catale et al., 2020; Lumertz et al., 2022). Microglia and astrocytes are increasingly recognized for their roles in synaptic plasticity and circuit functioning in the adult hippocampus (Allen and Barres, 2005; Ekdahl, 2012). Furthermore, chronic systemic neuroinflammation has been shown to impair spatial memory and LTP in adulthood, driven by elevated hippocampal levels of TNFα and IL1β primarily produced by microglia (Min et al., 2009; Liu et al., 2012; York et al., 2020; Hajipour et al., 2023). The complete normalization of both CD11b and TNFα mRNA expression following neonatal OT treatment points to potent anti-inflammatory effects of early OT that may represent a primary mechanism of neuroprotection. Accumulating evidence demonstrates that OT can reduce microglial activation and pro-inflammatory cytokine production across multiple brain regions (Yuan et al., 2016; Kamrani-Sharif et al., 2023), with perinatal carbetocin (an OTR agonist) attenuating microglial activation at transcriptional and cellular levels and providing long-term neuroprotection (Mairesse et al., 2019). The neonatal timing of OT administration appears particularly critical, potentially intercepting neuroimmune priming before consolidation. The lack of significant changes in GFAP transcription suggests a limited contribution of astrocyte number to the cognitive consequences of NMS. However, stress has recently been shown to induce remodeling of astrocyte morphology in an oxytocin-dependent manner in the amygdala, modulating neuronal responses (Baudon et al., 2025). Whether similar morphological remodeling occurs in the hippocampus following NMS remains to be determined, and neonatal OT injections may restore initial astrocytic morphology independently of changes in cell number. The trend toward OTR upregulation in NMS males may reflect homeostatic compensatory responses to disrupted maternal care and reduced endogenous OT signaling during early development (Turrigiano and Nelson, 2004; Gazzo et al., 2021; Gieré et al., 2023; Illouz et al., 2025). Supporting this hypothesis, exogenous OT administration during P2-P12 normalized OTR mRNA expression in adulthood.

The abolition of LTP at Schaffer collateral-CA1 synapses in NMS males suggests a mechanistic explanation for spatial memory deficits, as LTP at these synapses is essential for encoding spatial information (Morris and Frey, 1997; Martin et al., 2000). This selective loss of LTP without disruption of LTD points to a mechanistic dissociation downstream of NMDA receptors activation. The transcriptional upregulation of GAD65 and KCC2 raises the possibility of an inhibitory shunt selectively impairing LTP induction: increased GAD65 expression would enhance tonic GABAergic conductance at CA1 synapses (Song et al., 2011), while KCC2 upregulation is expected to shift the chloride reversal potential to more hyperpolarized values, increasing the driving force for chloride influx through GABA_A_ receptors (Rivera et al., 1999). Such changes would attenuate temporal summation of fEPSPs, thereby preventing the sustained depolarization required to relieve magnesium block of NMDA receptors (Bliss and Collingridge, 1993). LTD would nonetheless be preserved, as its induction requires only modest Ca²⁺ influx sufficient to activate calcineurin, rendering it less vulnerable to shunt-mediated attenuation of Ca²⁺ transients (Mulkey et al., 1994; Malenka and Bear, 2004; Li et al., 2012). Direct validation of this hypothesis through gramicidin perforated-patch recordings or chloride imaging would provide mechanistic validation.

In concordance with this hypothesis, alterations of inhibitory synaptic transmission mediated by GABA in the hippocampus have been revealed following neonatal stress and early inflammation (Veerawatananan et al., 2016; Liang et al., 2019; Aghighi Bidgoli et al., 2020; Gazzo et al., 2021). These alterations fit within a broader developmental context. Particularly, in most mature neurons, KCC2 activity maintains low intracellular chloride, enabling GABA_A_ receptor activation to hyperpolarize the membrane. During development, immature neurons express low KCC2 and high NKCC1, resulting in depolarizing GABA action (Ben-Ari, 2002). During postnatal maturation, a progressive increase in KCC2 shifts GABA toward its mature hyperpolarizing function (Rivera et al., 1999; Cordero-Erausquin et al., 2005; Kahle et al., 2015). Delayed hippocampal chloride shift following ELS has been previously documented (Veerawatananan et al., 2016), supporting the hypothesis that NMS disrupts GABAergic developmental trajectories. The overexpression of GAD65 and KCC2 mRNA in dorsal CA1 observed in the present study suggests alterations in GABAergic signaling and chloride homeostasis. In line with this, elevated KCC2 transcripts have already been reported in the dorsal CA1 of NMS males (Illouz *et al*., 2025). Adult male rats exposed to neonatal inflammation, or to perinatal ethanol, also show an increase in KCC2 protein in their dorsal CA1, associated with LTP deficits via enhanced GABAergic synaptic inhibition (Silvestre de Ferron et al., 2017; Rong et al., 2023).

Early-life OT treatment fully restored LTP while preserving LTD. Neonatal OT administration also markedly reduced GAD65, NKCC1 and KCC2 mRNA expression below control levels in P60 NMS+OT animals. The transcriptional downregulation of GAD65 in NMS+OT rats is consistent with reduced tonic GABAergic conductance (Song et al., 2011). Since OT also accelerates GABAergic maturation through OTR-mediated upregulation of KCC2 expression (Leonzino et al., 2016), this peptide facilitates the transition to mature hyperpolarizing transmission and optimal E/I balance during hippocampal development. Thus, exogenous OT may have modified the GABAergic developmental program, though whether this represents accelerated maturation, compensatory reorganization, or overcorrection requires further investigation with developmental time-course analyses and functional assessments of GABA polarity and chloride homeostasis. This reduction of GABAergic tone may act synergistically with OT-mediated facilitation of LTP through MAPK (mitogen-activated protein kinases), PI3K (phosphoinositide 3-kinases) and ERK (extracellular signal-regulated kinase) signaling cascades downstream of OTR activation (Tomizawa et al., 2003; Lin et al., 2012; Lin and Hsu, 2018). Literature supports this view, as OT application restores LTP in stressed animals across both dorsal and ventral CA1 (Lin et al., 2012; Lee et al., 2015), and adult intranasal OT administration to NMS rats also restores hippocampal LTP (Joushi et al., 2021). A parallel brain-derived neurotrophic factor (BDNF)-dependent mechanism may also contribute, given that NMS reduces hippocampal BDNF expression in adulthood (Aisa et al., 2009), BDNF signaling is a facilitator of LTP (Bramham and Messaoudi, 2005) and neonatal OT modulates BDNF signaling (Bukatova et al., 2023). These findings suggest that the OT intervention prevents the developmental emergence of aberrant GABAergic inhibition.

The paradoxical transcriptional upregulation of SYN and PSD95 in NMS males, despite previously reported protein-level reductions (Cui et al., 2020; Wang et al., 2020), may reflect a homeostatic scaling response to chronic synaptic under-activation consistent with the GABAergic shunt hypothesis (Turrigiano and Nelson, 2004), or post-transcriptional silencing via stress-induced miRNA upregulation (Melchior *et al*., 2022). The reduction of these transcripts below control levels in NMS+OT rats may reflect a reversal of the maladaptive homeostatic upregulation, consistent with a shift toward fewer but functionally competent synapses capable of sustaining LTP. Protein-level validation of all transcriptional findings reported here is important for future work. As OT was administered systemically, the relative contributions of direct central versus peripheral mechanisms to the observed hippocampal effects also remain to be determined.

These findings carry significant translational implications, as ELS is associated with reduced hippocampal volume during childhood or in adulthood (Teicher et al., 2016, 2018; Chau et al., 2019). Genetic variation in OTR signaling also interacts with early trauma exposure to predict hippocampal structural alterations in humans (Malhi et al., 2020), suggesting that endogenous oxytocinergic dysfunction may mediate stress-related hippocampal pathology across species. The present demonstration that neonatal OT administration prevents circuit-level dysfunction persisting into adulthood establishes the early postnatal period as a critical window during which oxytocinergic signaling shapes hippocampal maturation and confers lasting resilience. These findings raise the possibility that early oxytocinergic interventions could be explored in at-risk pediatric populations. Notably, existing clinical evidence that intranasal OT administration improves long-term memory and reduces anxiety in adults further supports the therapeutic relevance of oxytocinergic modulation, and the present findings extend this rationale to the neonatal period as a potentially more impactful window of intervention (Brambilla et al., 2016; Riem et al., 2020). Defining this neonatal window and identifying early biomarkers of oxytocinergic vulnerability will be essential to translate these neuroprotective mechanisms into targeted interventions for children exposed to early adversity.

## Acknowledgments

Research work has been financed by recurrent funding from the CNRS and the University of Strasbourg, as well as by grants from the Agence nationale de la recherche (ANR) as part of the Programme d’investissement d’Avenir (ANR-17-EURE-022 contract, EURIDOL), the Région grand est (ClueDOL, Fonds de coopération régionale et de recherche). PP is a senior member of the Institut Universitaire de France. HI and MJ received a fellowship from french Ministère chargé de l’enseignement supérieur et de la recherche.

