## Supplemental Table 1 for "Neonatal oxytocin prevents sex-specific spatial memory deficits induced by maternal separation through restoration of hippocampal synaptic plasticity in males"

Supplementary Table 1: Primer sequences used in this study.

| **Gene** | **Primers sequences (forward/reverse)** | **NCBI reference** |
| --- | --- | --- |
| **GAPDH** | GGCCTTCCGTGTTCCTAC  TGTCATCATACTTGGCAGGTT | NM017008 |
| **CD11b** | CTGCCTCAGGGATCCGTAAAG  CCTCTGCCTCAGGAATGACATC | NM012711 |
| **TNFα** | GGGCTGTACCTTATCTACTCC  TATGAAATGGCAAATCGG | NM012675 |
| **GFAP** | TGGAGAGAATTGAATCGC  CCGGAGTTCTCGAACTTCCTC | NM010277 |
| **GAD65** | GACCTGTCCTATGACACGG  AGCTTCAAATCCAGTAGTCCC | NM012563.2 |
| **NKCC1** | GGGCCTCCTCACACGAAGAA  TGAGGAGCCGAGGGTACTTCA | NM019229 |
| **KCC2** | ACTACAGCTGGCCACCTCGC  ATGCTGCCCTCAGAGAAACGC | NM134363 |
| **Synaptophysin** | CTTTGCCATCTTCGCCTTT  ATGTTGAGGGCACTCTCCGTC | BC099798-1 |
| **PSD95** | CATCATCATCCTTGGACCCAC  CGCTTAGGACGTGCTGTATG | NH018621.2 |
| **OTR** | CCCGTGAAGAGCATGTAGATC  CAAAGAAGCTTCTGCCTTCAT | NM012871 |
